# Primate Saccade Rhythmicity

**DOI:** 10.1101/2023.09.27.559710

**Authors:** Tim Näher, Yufeng Zhang, Silvia Spadacenta, Martina Pandinelli, Lisa Bastian, Peter Thier, Ziad M. Hafed, Pascal Fries

## Abstract

Rhythmic sampling is a hallmark of many sensory systems across diverse organisms. In primate vision, rapid foveating eye movements repetitively sample the visual environment at a rate of approximately 4 Hz. Yet, the observed inter-saccadic interval distributions might equally well be produced by a non-rhythmic stochastic process as by a —putatively variable— rhythmic generator. This raises the fundamental question whether primate visual sampling differs from the numerous examples of rhythmic sampling strategies exhibited across sensory modalities and species. Here, using experiments in humans, macaques, and a marmoset, as well as statistical approaches adopted from studies of temporal structure in neuronal spiking patterns, we show that primate saccade generation is unambiguously rooted in a rhythmic source. This finding was remarkably consistent across the three primate species despite their significant evolutionary distance. Importantly, we find that saccade rate undergoes smooth, slow fluctuations on the scale of seconds, and accounting for those rate fluctuations was crucial for revealing the true degree of saccade rhythmicity. Thus, exact saccade timing is determined by an interaction between the slow saccade-rate fluctuations and the more temporally-local rhythmic generator. This demonstration of rhythmicity in overt oculomotor sampling behavior provides a link to the fundamentally rhythmic nature of many central perceptual and cognitive processes, and places primate saccadic sampling into the large family of rhythmic active sampling behaviors.

**Significance Statement:** Cognition and perception are shaped by brain rhythms, and rhythmic sampling governs covert attention. Yet, whether overt attentional sampling by saccades, the most common primate visual exploration behavior, is also rhythmic, has remained elusive. Here, we present a variety of artificial and natural images to humans, macaques, and a marmoset, and record their eye movements during free exploration. We find that saccade rate fluctuates widely and slowly across seconds, and that accounting for those rate fluctuations reveals a pronounced temporally local saccade rhythmicity. This demonstration of rhythmicity in primate saccadic sampling reunites overt and covert attention under a common rhythmic principle and places primate visual exploration alongside rodent whisking and sniffing as a rhythmic active sensing behavior.

## Introduction

Vision is an active process: rather than passively receiving stimuli, organisms actively reposition their eyes to selectively sample behaviorally relevant information from the environment. In primates, saccadic eye movements shift the fovea to successive locations, creating a sequence of high-acuity “snapshots”. During natural viewing, saccades occur repetitively, with inter-saccadic intervals (ISIs) in the range of few hundred milliseconds, yielding mean saccade rates of 2–6 Hz^1^. This repetitive sampling bears a striking resemblance to other active-sensing behaviors such as whisking and sniffing in rodents ^2^. Those behaviors are known to be strongly rhythmic, raising the question of whether similar intrinsic rhythmic processes also govern saccade generation^3^.

Several influential accounts of saccade behavior have, in fact, implicitly contained or predicted some degree of rhythmicity in the saccadic system. Classical descriptions of the temporal structure of saccades emphasized two temporal regularities: 1) a refractory period following each saccade, during which the probability of initiating the next movement is suppressed, and 2) a unimodal shape of the ISI distribution^4^. Both of these saccade characteristics can be explained by a saccade generation model based on a repetitive rise-to-threshold accumulation. Together, this implies that saccades show distinct temporal structure.

However, detailed inspection of saccade timing patterns reveals large variability in observed ISIs across many behavioral tasks, resulting in heavily skewed ISI distributions with substantial tails produced by fairly long ISI values. Such skew significantly complicates the interpretation of saccade timing as a rhythmic process, with some accounts even going as far as arguing that skewed ISI distributions are a manifestation of a non-rhythmic stochastic process governing saccade generation ^5^. This, if true, would challenge prevailing assumptions about the strong links between eye movements and cognitive processes like attention^6–9^ and visual sensitivity^10,11^, which are themselves believed to be rhythmic. Therefore, demonstrating and understanding the properties of saccade-system rhythmicity is a fundamental problem in primate systems neuroscience.

In this work, inspired by statistical analysis approaches historically used to study temporal dynamics in neuronal spiking, we demonstrate genuine saccade-system rhythmicity in the primate brain. Using experiments with humans, macaques and a marmoset performing the same visual tasks, we show that the observed highly skewed ISI distributions arise from a rhythmic generator being modulated by slow saccade-rate fluctuations, putatively due to factors like arousal, scene statistics, or task context. When we correct for slow saccade-rate fluctuations, we find that rhythmicity shows a genuine positive relation to saccade rate. These findings reconcile previous conflicting accounts of saccade timing and demonstrate that a temporally local rhythmic generator does exist for saccades. Beyond providing a link between rhythmicity in covert attentional sampling and overt saccadic sampling, our results motivate investigation into the neuronal mechanisms underlying rhythmic saccade drive.

## Methods Summary

We studied 22 human participants, two rhesus macaques, and one common marmoset, while they visually foraged on regular or irregular target arrays or natural images. Human subjects additionally viewed a naturalistic movie that was largely free of cuts. All procedures were approved by the responsible ethics committee or authorities. Eye movements were recorded with eye-tracking systems, and analyzed using species-specific convolutional neural networks for saccade detection, accepting only saccades detected simultaneously in both eyes. We quantified saccade rhythmicity using (1) a gamma renewal model fitted to inter-saccadic intervals (ISIs), estimating its shape parameter from within-trial normalized ISIs using a Dirichlet likelihood, and (2) the rate-corrected rhythmicity index (RCRI), based on variability between adjacent ISIs. To characterize slow saccade-rate fluctuations and their effects on rhythmicity measures, we analyzed trial-wise and within-trial saccade rates, ISI serial correlations, and simulations of stationary and rate-varying gamma renewal processes. Statistical analyses included species-specific regression, comparisons across saccade-rate bins, and tests of differences in rhythmicity, with significance assessed at α = 0.05 unless otherwise specified. Details of the methods can be found in SI Appendix.

## Results

We characterized the nature of saccade rhythmicity in three primate species by having them view static or dynamic scenes. In each case, we assessed to what extent inter-saccadic intervals (ISIs) could be thought of as being generated by a rhythmic process, and we did so by relating ISIs to slower saccade-rate fluctuations. We hypothesized that ISI distributions can be effectively modeled as a doubly-stochastic process, with one component governed largely by a relatively regular oscillator, and another component that slowly alters the period of the oscillator and thus the overall saccade rate. In what follows, we first show that saccade rate substantially fluctuates in all three species, and we then explain how accounting for such fluctuations reveals the degree of intrinsic rhythmicity in saccades.

### Saccade rates vary across trials

We first used three different static displays (Figure 1a-c; see Methods for details of the displays and tasks). In displays of REGULAR (Figure 1a) or IRREGULAR (Figure 1b) dot arrays, a random subset of 20% of the dots were targets associated with reward when fixated. Similarly, in displays of natural IMAGEs (Figure 1c), after a random delay of 100 ms to 1900 ms and at a random location, a small white dot appeared that was a target associated with reward when fixated. The maximal time until target fixation was 5000 ms. This was sufficient for target detection in most cases, precluding a meaningful analysis of a potential relation between saccade rate and target-detection performance. Yet, an analysis of the relation between target-detection latency and saccade rate was possible. Higher saccade rates showed a modest relation with shorter latencies in humans (β = −0.19 on log latencies, R² = 0.11, n = 22 participants, performing a total of 3,959 trials). The two macaques showed the same relation (β = −0.35 on log latencies, R² = 0.22, n = 3,370 trials). For the marmoset, a software error prevented recording of the end-of-trial marker, so target-detection latencies were not available.

**Figure 1:**
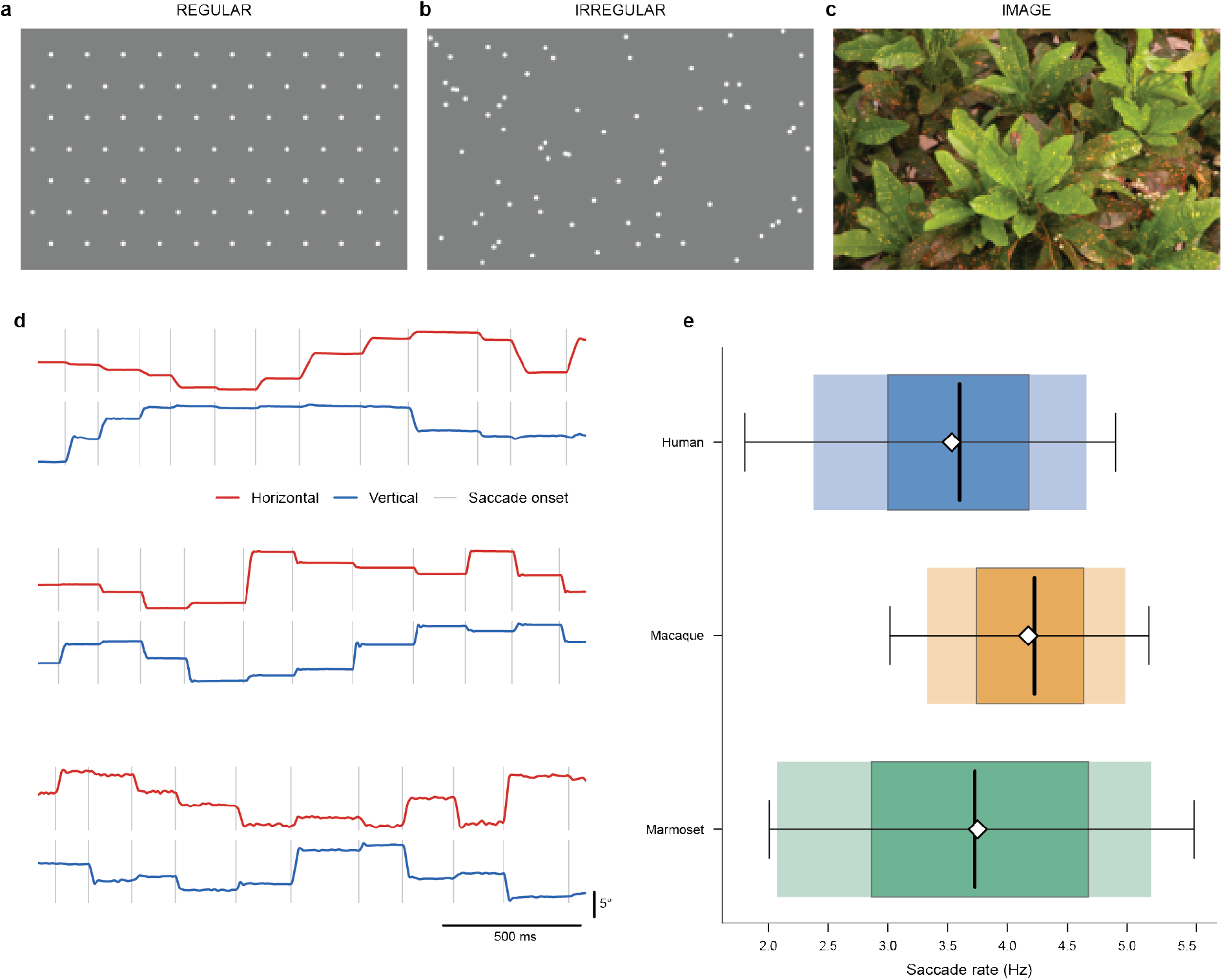
Saccade rates vary across trials. (a-c) Example task screens of the three conditions: REGULAR, IRREGULAR, and IMAGE. These conditions were used for all analyses in this figure. **(d)** Example eye traces from all three species, ordered as in **e**. Detected saccade onsets are indicated by vertical lines. Temporal scale bar indicates 500ms, vertical scale bar 5 degree of visual angle. **(e)** Box plots reflecting the distributions of saccade rates over trials. Trials were pooled over conditions (REGULAR, IRREGULAR, and IMAGE) and subjects, separately for the three species. Median [interquartile] rates were 3.60 [3.00, 4.18] Hz (human, n = 6929 trials), 4.22 [3.74, 4.64] Hz (macaque, n = 6057 trials), and 3.73 [2.86, 4.68] Hz (marmoset, n = 568 trials). The 5th–95th percentile ranges were 1.80–4.90 Hz, 3.02–5.18 Hz, and 2.01–5.56 Hz, respectively. The darker shaded box spans the interquartile range (25th–75th percentile), the lighter shaded region extends from the 10th to 90th percentile, the vertical black line marks the median, the white diamond indicates the mean, and the whiskers with caps denote the 5th–95th percentile range. Mean ± SD of the saccade rates were 3.53 ± 0.95 Hz in humans, 4.17 ± 0.67 Hz in macaques, and 3.75 ± 1.18 Hz in the marmoset. Saccade rate was computed per trial as the number of saccades divided by trial duration (s), after excluding the first 100 ms following array/image onset to minimize onset-transient effects.

For testing our abovementioned hypothesis, we quantified the variability in saccade rates across trials for the three investigated species: human, macaque and marmoset (Figure 1d). For each species, saccade rates varied widely rather than clustering tightly around a single characteristic value (Figure 1e). In humans, trial-wise rates spanned more than 3 Hz across the central 90% of trials (5th–95th percentile: 1.80–4.90; mean ± SD: 3.53 ± 0.95 Hz), indicating substantial variability. Macaques showed a similarly broad distribution (5th–95th percentile: 3.02–5.18, mean ± SD: 4.17 ± 0.67 Hz), while the marmoset exhibited the largest relative spread (5th–95th percentile: 2.01–5.56, mean ± SD: 3.75 ± 1.18 Hz). Thus, in all three species, saccade rates varied substantially across trials, indicating that the underlying process cannot be modeled with a single stationary rate.

### Gamma renewal models can reproduce heavy-tailed ISI distributions

The large variability in saccade rates observed in all tested species (Figure 1) has two important implications for the interpretation of saccade timing statistics. First, the empirically observed heavy-tailed ISI distribution reflects a marginal distribution aggregated over time, rather than the interval statistics generated at any fixed moment. When saccade rate varies, the overall ISI distribution becomes a mixture of many locally generated interval distributions with different means, such that heavy tails might arise even if saccades are generated by a locally rhythmic process. Second, rate fluctuations can introduce higher-order temporal structure that can affect spectral analyses of saccade rhythmicity.

To test these implications explicitly, we modeled saccade generation as a doubly stochastic gamma renewal process, with temporally local rhythmicity (reflected by the shape parameter *k*) that remained constant over trials, and a saccade rate (reflected by the scale parameter *θ_t_*) that fluctuated slowly across trials. In estimating *k*, we eliminated the rate component analytically, by transforming the ISIs of a given trial into proportions of their sum and fitting these proportions with a Dirichlet distribution (see Methods for details). The resulting *k* values are shown in Figure 2a, separately for all participants from all three species. In humans, fitted k values were 9.54 ± 1.78 (mean ± SD; 5th–95th percentile: 7.42–12.43; min–max: 7.36–15.51, n=22). Macaques showed similar values (monkey G: 10.0, monkey C: 14.6). The marmoset data yielded a gamma shape parameter of 24.06.

**Figure 2:**
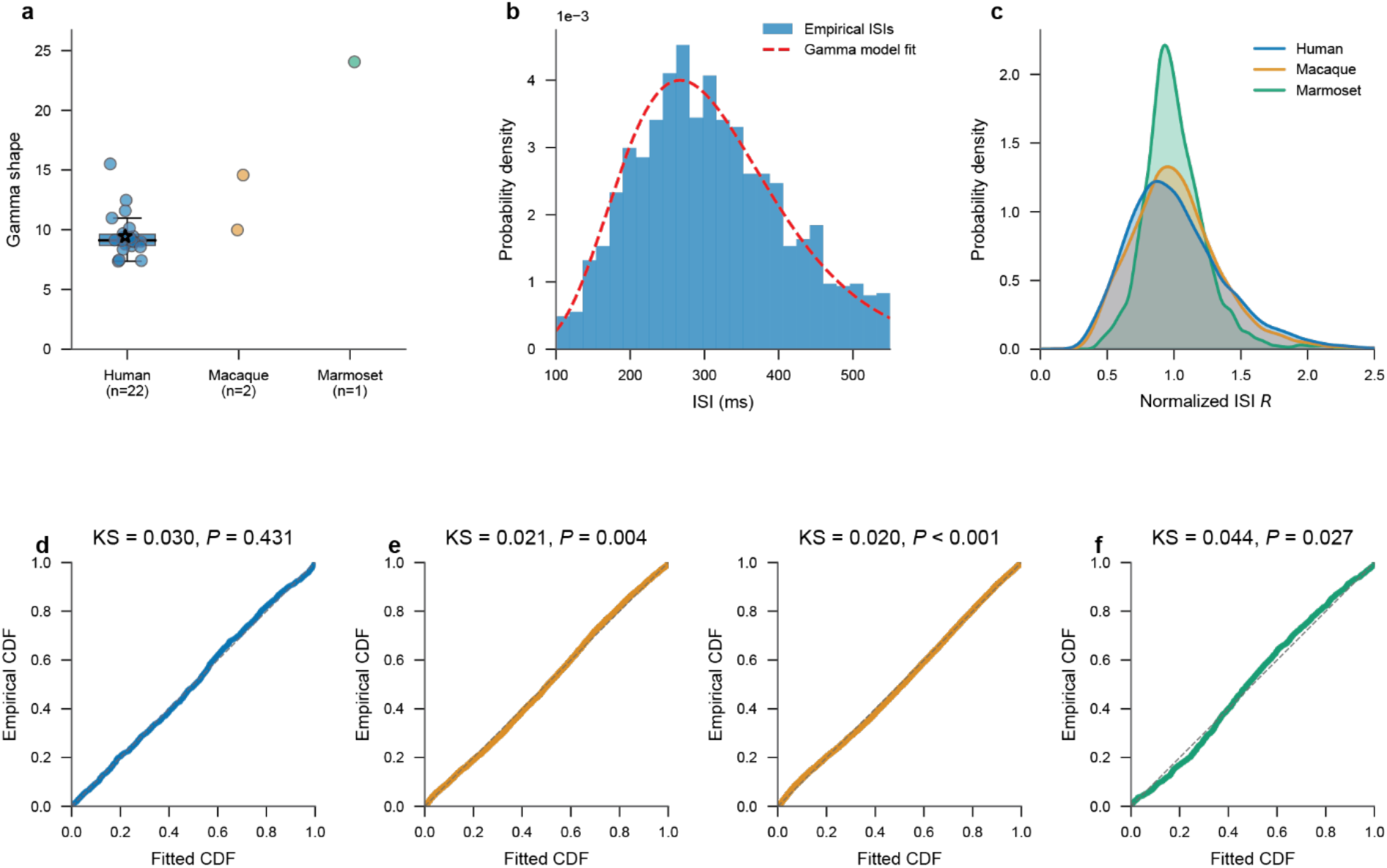
Gamma renewal models can reproduce heavy-tailed ISI distributions. Per participant, ISIs were modeled as a gamma renewal process with a common shape parameter k and a trial-specific rate. The rate was eliminated analytically by expressing each trial’s ISIs as proportions of their sum, which follow a Dirichlet distribution with the same k. k was the only free parameter and was estimated by maximum likelihood across all trials (see Methods). **(a)** Each point shows the shape parameter k for one individual participant; the star indicates the example human participant in panel **b**. The large k values indicate strong local rhythmicity: a gamma distribution with k ≈ 10 has a coefficient of variation of 0.32 and is near-symmetric. **(b)** Blue: Empirical ISI distribution from the example human participant. Red: ISI distribution predicted by the fitted model, computed as the average across trials of gamma densities that share the fitted k, each scaled by that trial’s own rate (its number of retained ISIs divided by their summed duration). The model reproduces the heavy-tailed ISI distribution despite being locally rhythmic with k close to ten, showing that the heavy tail arises from mixing across trial rates rather than from irregular local timing. **(c)** Distributions of ISIs normalized by the average of all other ISIs in the same trial, separately for all three species. **(d)** Example human probability-probability (PP) plot of the probability integral transform of the interval proportions against the uniform distribution, showing that the observed intervals follow the proposed gamma renewal model. For humans, all KS-tests were non-significant, indicating that the gamma model is a good fit. **(e)** same as **d**, but for the two macaques. **(f)** same as in **e**, but for the marmoset. Note that the KS tests were significant for the macaques and the marmoset.

Figure 2b shows the ISI distribution of an example human participant, together with the gamma model fit. The goodness of fit was assessed visually with probability-probability (PP) plots and statistically with one-sample Kolmogorov–Smirnov (KS) tests (Figure 2d-f).

In none of the human participants, the gamma distribution deviated significantly from the observed distribution (mean KS = 0.031, mean P = 0.590 with 0/22 rejected at α = 0.05), indicating that the gamma distributions provided an excellent fit (Figure 2d).

In macaques (Figure 2e), deviations were significant but small for both animals (monkey G: k = 10.0, KS = 0.020, P < 0.001, maximal deviation between cumulative densities: 2.0%; monkey C: k = 14.6, KS = 0.021, P = 0.004, maximal deviation between cumulative densities: 2.1%). Note that the tests were likely more sensitive in macaques than humans, because the number of retained ISIs was 10,930 for monkey G and 7,002 for monkey C and thereby much larger than the median ≈900 for the human participants. When we subsampled the number of macaque saccades to 900, deviations were significant in fewer than 2% of draws for either animal (1.7% for monkey C, 1.2% for monkey G). Note that the ISI distributions did not discriminate between comparable two-parameter families: fitted with a common shape and a trial-specific scale, and accounting for the 100–600 ms observation window, the gamma, lognormal, inverse Gaussian and Weibull distributions differed only negligibly in Akaike information criterion (median ΔAIC ≤ 8 across subjects). We retain the gamma because it is the renewal family whose interval proportions follow a Dirichlet distribution, which allows the saccade rate to be eliminated analytically rather than estimated. When we inspected the empirical and fitted distributions in detail, we found that in monkey G (k = 10.0), the deviation was highly significant but negligible in magnitude, reflecting that animal’s large sample: quantile ranges matched the model closely (all ratios of empirical to model quantile ranges were between 0.98 and 1.00); In monkey C (k = 14.6), the central body was more concentrated than predicted (empirical IQR 8% narrower than the model IQR), while the extreme tails were wider (1st–99th percentile range 8% wider), and kurtosis exceeded the model prediction (empirical: 3.39, model: 0.95), indicating a leptokurtic distribution, i.e., a distribution with a sharper peak and heavier tails than the fitted gamma renewal model.

The marmoset (Figure 2f) showed a similar pattern as monkey C, but even more pronounced (k = 24.06, KS = 0.044, P = 0.027, maximal deviation between cumulative densities: 4.4%). When we subsampled the 1,103 retained marmoset ISIs to 900, deviations remained significant in 36% of draws. Unlike in the macaques, the lognormal distribution fitted marginally better than the gamma (ΔAIC = −12.7), whereas the gamma clearly outperformed the Weibull (ΔAIC = +12.4) and the inverse Gaussian (ΔAIC = +71.0). The central body was even more concentrated (IQR 14% narrower), and the extreme tails even wider (1st–99th range 28% wider, excess kurtosis empirical: 7.91, model: 0.64), than in monkey C, indicating an even more pronounced leptokurtic distribution.

The leptokurtic empirical distributions of normalized ISIs in monkey C and the marmoset suggest a highly rhythmic core saccade generation process plus few very long ISIs. Those very long ISIs might be due to the saccade-detection procedure missing a few saccades.

To visualize the distributions of rate-corrected ISIs, we divided, per trial, each ISI by the average of all other ISIs, i.e., by the expected ISI given the saccade rate, to arrive at rate-normalized ISIs (see Methods for details). The normalized ISIs were pooled over trials, conditions and participants, separately per species, and are shown in Figure 2C.

Having established a substantial rhythmic component in primate saccade timing, we next asked how slow rate fluctuations affect two previously used metrics of event rhythmicity or variability, respectively. Amit et al.^5^ had used spectral measures to quantify saccade rhythmicity. Holt et al.^12^ had used a local estimate of the coefficient of variation (CV) to quantify the variability across neuronal inter-spike intervals. The conventional CV is the ratio of the standard deviation to the mean across all intervals, and Holt et al. note that it is an inaccurate metric if rates change. Therefore, they introduce C_V2_, which avoids sensitivity to changing rates by comparing only adjacent intervals^12^ (see Methods for details). We compared (1) spectral peak power and, (2) the rate-corrected rhythmicity index, *RCRI*. The RCRI is a monotonic reparameterization of the well-established C_V2_ metric^12^, such that RCRI increases with saccade rhythmicity (see Methods for details). We simulated stationary and non-stationary gamma renewal processes that had similar local rhythmicity and similar mean rates yet differed with regard to the presence or absence of rate fluctuations over time (Figure 3).

**Figure 3:**
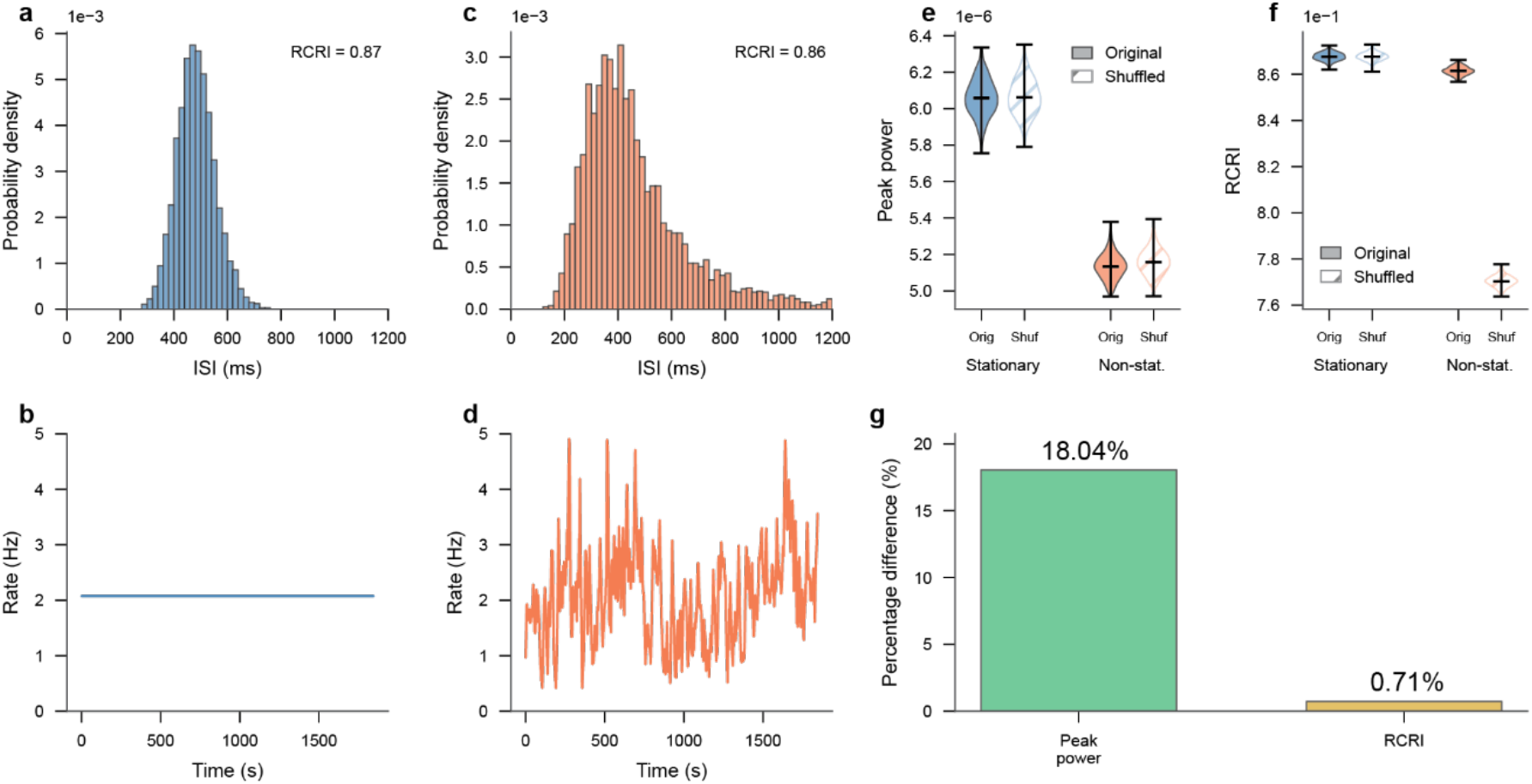
**Slow rate fluctuations confound spectral rhythmicity measures**. All panels show the results of simulations of gamma renewal processes with a fixed shape parameter of 44.4. **(a, b)** Example stationary gamma renewal process with fixed rate (b) that is based on the mean of the time series in (d). The resulting marginal ISI distribution (a) is narrow and devoid of a heavy tail. Note that the x-axis scale in panel (a) is in milliseconds, and in panel (b) is in seconds. (**c, d)** Example non-stationary gamma renewal process with an empirical rate time series (d) from a free-viewing trial of the MOVIE condition. The resulting marginal ISI distribution (c) is heavy-tailed despite being based on the same shape parameter as panel (a) and thereby the same local rhythmicity. Note that the x-axis scale in panel (c) is in milliseconds, and in panel (d) is in seconds. **(e)** Peak power (2–6 Hz) across 500 simulations, comparing original and ISI-shuffled saccade trains for both stationary and non-stationary gamma renewal processes. **(f)** Rate-corrected rhythmicity index (RCRI) for the same original and ISI-shuffled stationary and non-stationary processes across 500 simulations. Note that the y-axis is highly zoomed in and reveals that the RCRI is highly similar between the stationary and non-stationary process, that shuffling reduced RCRI only for the non-stationary process, whereas RCRI of the stationary process was shuffle-invariant, as required for shuffling to be a valid control. **(g)** Percent difference between the same stationary and non-stationary processes in peak power (left) and RCRI (right).

Figure 3a,b shows an example simulated stationary gamma renewal process with a shape parameter of 44.4 and a constant rate corresponding to the mean rate of the empirical rate time course shown in Figure 3d. This process produced a narrow ISI distribution (Figure 3a), and an RCRI of 0.87, reflecting its strong local rhythmicity. Figure 3c,d shows an example simulated non-stationary gamma renewal process with the same shape parameter of 44.4, but a rate that followed a time course empirically observed during a long free-viewing MOVIE-condition trial. Despite producing a heavy-tailed ISI distribution (Figure 3c), the RCRI of 0.86 was nearly identical to the stationary process, indicating very similar local rhythmicity once slow rate fluctuations on the time scale of seconds were accounted for.

Repeating this simulation 500 times showed that spectral measures of rhythmicity, such as peak power^5^, are strongly affected by rate changes (Figure 3e), while the RCRI is hardly affected (Figure 3f, note the highly zoomed-in y-axis). The difference between the stationary and non-stationary condition is 18.04% for peak power, while it is merely 0.71% for the RCRI (Figure 3g).

An ISI-shuffled control within each condition (lighter violins in Figure 3e,f) revealed that peak power was unaffected by shuffling regardless of stationarity, demonstrating that spectral measures alone do not validly capture local rhythmicity in the face of slow rate fluctuations. In contrast, RCRI dropped after shuffling selectively in the non-stationary condition while remaining invariant for the stationary process, confirming its sensitivity to the local serial dependencies imposed by rate modulation.

Together, Figures 2 and 3 show that two properties of the temporal dynamics of saccade generation, namely heavy-tailed ISI distributions and attenuated spectral peaks, can both arise from a locally rhythmic saccade generator operating under a slowly varying rate. These slow rate fluctuations will be characterized in more detail later. In this framework, local rhythmicity of successive intervals is preserved, while slow rate fluctuations shape global interval statistics, such as the marginal ISI distribution, and introduce higher-order temporal structure that degrades standard spectral measures of rhythmicity. Having disentangled these rhythmic and rate-dependent components, we next used the RCRI to isolate the rhythmic component and compare it across tasks and species.

### Saccade rate explains across-task differences in rhythmicity

So far, we have established that slow rate fluctuations shape ISI distributions and degrade spectral measures of saccade rhythmicity, and that the RCRI provides a robust estimate of local rhythmicity. Because the RCRI is nearly immune to slow rate variability, we next investigated whether RCRI shows a genuine physiological relation to saccade rate. We calculated RCRI for humans, macaques, and marmoset performing a visual foraging task in three conditions: a regular hexagonal grid (REGULAR), a spatially jittered version of the same grid (IRREGULAR), and a natural-image condition (IMAGE). The REGULAR and IRREGULAR conditions produced visual inputs with systematically varying spatial periodicity, while the naturalistic IMAGE condition further enhanced ecological validity.

Figure 4a shows average saccade rates across conditions for each species. In all species, saccade rates varied substantially between conditions, with the IMAGE condition generally eliciting lower saccade rates in humans and macaques, yet higher rates in the marmoset. Thus, task context robustly modulated saccade rate in all three species.

**Figure 4:**
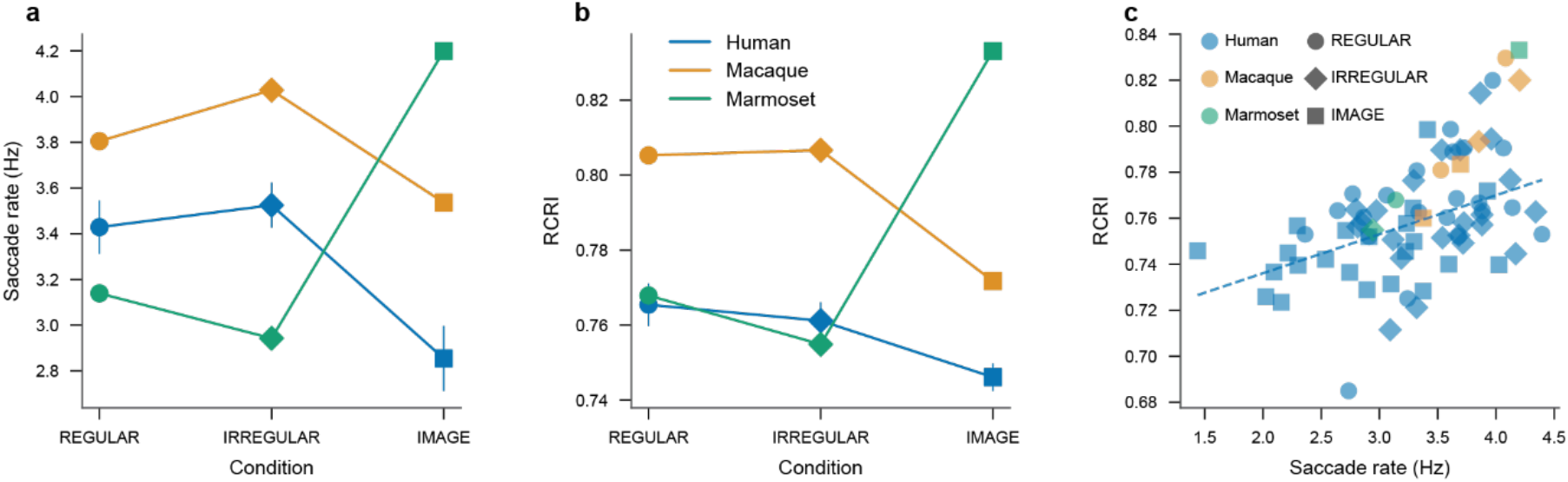
Saccade rates and RCRI across tasks and species. **(a)** Mean saccade rate across task conditions (REGULAR, IRREGULAR, IMAGE) for humans, macaques, and marmoset. Task context robustly modulates saccade rate in all species. **(b)** RCRI across the same conditions, showing condition-dependent changes that closely mirror saccade rates. **(c)** Relationship between saccade rate and RCRI for human data (blue symbols). Each symbol corresponds to the combination of a human participant and a task condition. Human data show a significant positive correlation (Spearman ρ = 0.47, P = 6.4 × 10⁻⁵; dashed line shows the least-squares fit, based on human data only). Macaque and marmoset data were not included in the correlation analysis yet fall along the same trend. Importantly, RCRI is invariant to uniform scaling of ISIs (see Methods), meaning the positive correlation between rate and rhythmicity cannot arise from a trivial dependence of the RCRI metric on ISI duration.

Figure 4b shows the corresponding RCRI values. Within each species, RCRI varied systematically with condition in a manner closely mirroring changes in saccade rate. Conditions associated with higher saccade rates also exhibited higher RCRI, whereas conditions with lower saccade rates showed reduced RCRI. This shared modulation suggests a genuine physiological relation between saccade rate and local saccade rhythmicity.

To quantify this relationship, we examined the association between saccade rate and RCRI. Figure 4c shows this relationship for human participants, with each dot corresponding to one of the tasks in one participant. This revealed a positive correlation between saccade rate and RCRI (Spearman ρ = 0.47, P = 6.4 × 10⁻⁵). A sensitivity analysis of how this correlation depends on analysis-parameter choices is shown in *SI appendix*, Fig. S1B. The corresponding data points from the two macaques and the marmoset fall into the human point cloud, descriptively pointing towards a similar relationship across species despite differences in absolute rates.

To disentangle the effects of saccade rate and task condition, we fit a linear model predicting RCRI from average saccade rate and condition. In humans, saccade rate was a significant predictor of RCRI (β = 0.0142 per Hz, P = 0.0041), whereas neither the IRREGULAR nor IMAGE condition differed from REGULAR once rate was accounted for. Thus, after controlling for saccade rate, task condition no longer explained additional variance in rhythmicity. Similar patterns were observed in macaques, where rate showed a strong positive effect on RCRI (β = 0.082 per Hz, P = 0.0030), while condition effects were not significant. The marmoset data followed the same trend descriptively, although formal inference was not possible due to the limited sample size.

We also analyzed whether RCRI was related to target-detection latency. In humans, higher RCRI values showed a weak relation with shorter latencies (β = −0.46 on log latencies, R² = 0.006, n = 22 participants, performing a total of 3,959 trials). In the two macaques, the corresponding relation was stronger (β = −1.28 on log latencies, R² = 0.04, n = 3,370 trials). Yet, these relations might be due to the relation between RCRI and saccade rate, together with the relation between saccade rate and latency reported above. Indeed, in humans, when saccade rate was accounted for, RCRI showed no relation to target-detection latency (β = −0.04, 95% CI [−0.22, +0.15], *P* = 0.70; adding RCRI left the explained variance unchanged). In the two macaques, a relation remained after accounting for saccade rate (β = −1.00, 95% CI [−1.21, −0.79], *P* < 0.001), although it explained little additional variance (ΔR² = 0.019) and, with two animals, applies to this sample rather than to the population.

Together, these results demonstrate differences in RCRI across tasks and species. Intriguingly, while the RCRI metric corrects for slow rate fluctuations, the RCRI differences could be accounted for by differences in saccade rate. This suggests a genuine physiological relation between saccade rate and saccade rhythmicity.

### Saccade rhythmicity as function of saccade rate

As an independent and nonparametric validation of the relationship between saccade rate and rhythmicity, we next binned trials by their mean saccade rate and assessed rhythmicity within these rate-defined trial subsets. For each species separately, trials were assigned to tertiles of saccade rate (low, medium, high) using quantile binning. Within each bin, we assessed rhythmicity with autocorrelation functions and the RCRI. Visual inspection of the autocorrelation functions suggested that across humans, macaques, and marmoset, higher-rate trials consistently exhibited stronger rhythmicity (Figure 5a,d,g): In all species, rhythmicity as indexed by the side peaks was clearest for the high-rate bin and progressively weakened in the medium-and low-rate bins. Correspondingly, ISI distributions for increasing saccade rates shifted systematically toward narrower distributions (Figure 5b,e,h). Most importantly, RCRI showed robust, monotonic increases with saccade rate (Figure 5c,f,i). In humans, RCRI differed between all pairs of rate bins (high vs. medium: *t* = 8.712, *P* = 4.38 × 10⁻¹⁸; high vs. low: *t* = 19.299*, P* = 1.16 × 10⁻⁷⁷ medium vs. low: *t* = 11.474, *P* = 8.49 × 10⁻³⁰). The same pattern was observed in macaques (high vs. medium: *z* = 5.877, *P* = 4.18 × 10⁻⁹; high vs. low: z = 12.860, *P* = 7.58 × 10⁻³⁸; medium vs. low: *z* = 7.777, *P* = 7.41 × 10⁻¹⁵) and in the marmoset (high vs. medium: z = 8.654, *P* = 4.97 × 10⁻¹⁸; high vs. low: *z* = 9.165, *P* = 4.95 × 10⁻²⁰; medium vs. low: *z* = 5.676, *P* = 1.38 × 10⁻⁸).

**Figure 5.**
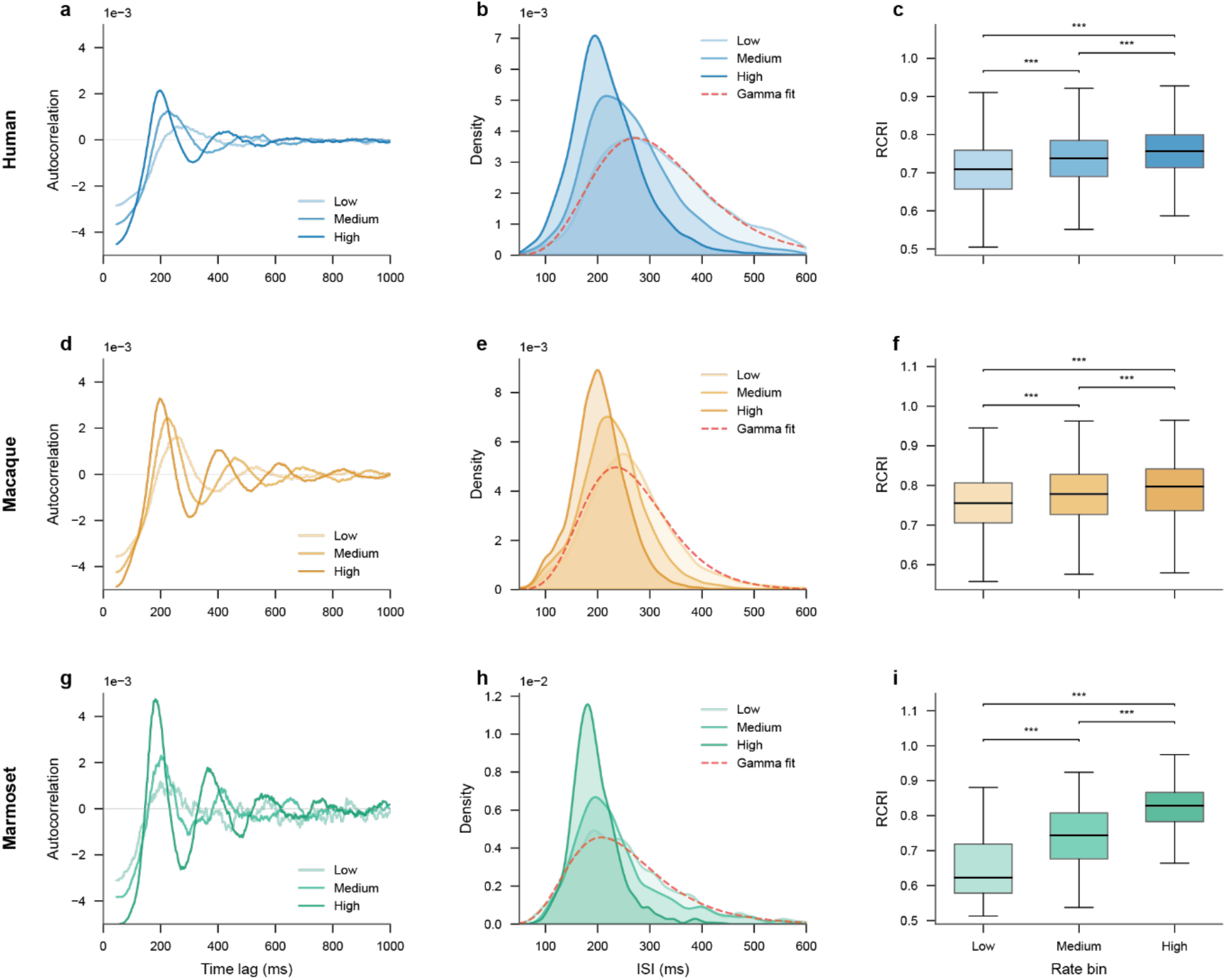
Rhythmic structure strengthens monotonically with saccade rate. (a, d, **g)** Autocorrelation functions of saccades for trials binned into low, medium, and high saccade-rate tertiles, shown separately for humans, macaques, and marmoset. Higher-rate trials exhibited more pronounced oscillatory side peaks. **(b, e, h)** ISI distributions for the same rate bins. Increasing saccade rate is associated with shorter ISIs and narrower ISI distributions. We repeated the analysis of Figure 2b for the lowest saccade rate bin, and plotted the ISI distributions approximated via doubly stochastic gamma fit as red dashed lines. **(c, f, i)** RCRI across rate bins for each species. RCRI increases monotonically from low-to high-rate bins.

This rate-binned analysis provides converging evidence that saccade rhythmicity increases monotonically with saccade rate for each of the investigated primate species. This conclusion is consistently supported by the side peaks in the autocorrelation function, by the shapes of the ISI distributions, and finally by the RCRI. Thus, the observed relationship between saccade rate and rhythmicity is not an artifact of a specific metric, modelling assumption, or analysis framework, but reflects a robust property of saccade rhythmicity across species.

### Slow rate fluctuations induce higher-order temporal dependencies

Having established that saccade rate varies substantially across trials (Figure 1), we next investigated saccade-rate dynamics during free viewing of the MOVIE condition in ≈30 min long uninterrupted trials. Figure 6a shows in gray an example saccade-rate time course from a single free-viewing trial, calculated on the basis of consecutive 3 s windows. Although this time course was noisy, a pronounced component fluctuating on the timescale of tens to hundreds of seconds was evident. We quantified this temporal structure in the saccade-rate time course (based on 3 s windows) by computing the autocorrelation function (Figure 6b). The autocorrelation was positive for short lags and decayed gradually with increasing lag, suggesting slow temporal structure. This effect did not arise trivially from calculating rate in finite windows as shown by a Monte Carlo null model (Figure 6b orange region; see also Methods).

**Figure 6.**
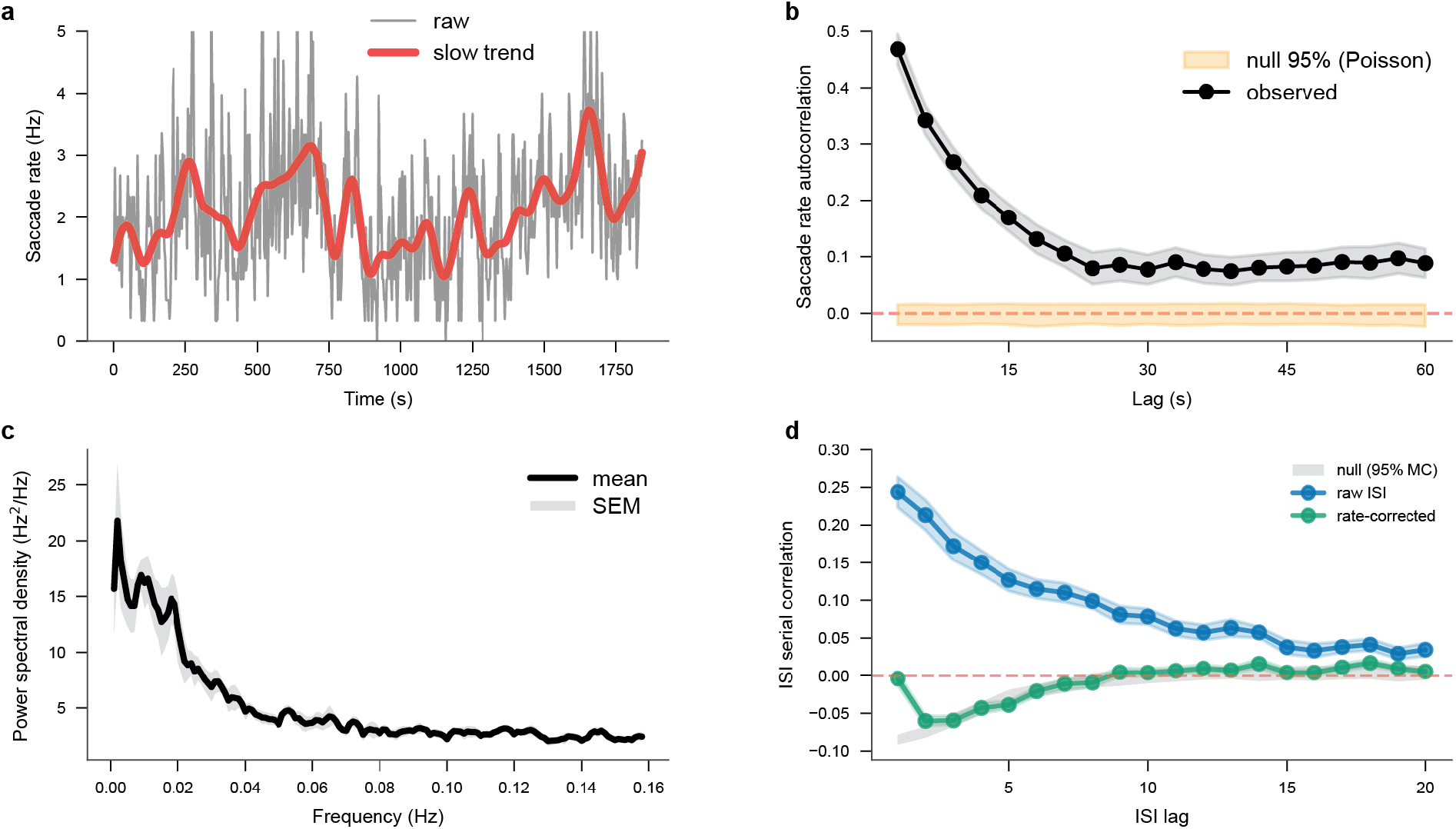
**Slow rate fluctuations generate higher-order temporal dependencies**. **(a)** Example saccade-rate time course from a single free-viewing MOVIE-condition trial. Gray line shows raw saccade rates for non-overlapping 3 s windows; red line shows slow rate fluctuations over tens to hundreds of seconds. Slow rate fluctuations were extracted by low-pass filtering the windowed rate series with a third-order zero phase Butterworth filter. Note that the x-axis scale in panel a is in seconds. **(b)** Autocorrelation of windowed saccade rate (black, mean ± SEM across participants), compared to a Monte Carlo null envelope derived from stationary homogeneous Poisson processes (orange, 95% CI). Non-overlapping 3-second windows. **(c)** Power spectral density of windowed saccade rate (mean ± SEM across participants), showing concentration of power at very low frequencies, consistent with slow rate fluctuations. **(d)** Serial correlation of raw ISIs (blue, mean ± SEM across participants) and rate-corrected ISIs (green, mean ± SEM across participants), against a Monte Carlo null band (grey, 95%) obtained by passing surrogates with no dependency beyond rate drift through the identical rescaling pipeline.

Spectral analysis of the rate time course further supported the presence of slow rate fluctuations. The power spectral density of the windowed rate estimates was dominated by very low frequencies and showed a ∼1/f-like decay, with no distinct spectral peak (Figure 6c).

We investigated whether these slow rate fluctuations induced higher-order temporal dependencies in saccade timing. We computed the serial correlation of ISIs excluding intervals overlapping blinks. The serial correlation was positive for short lags and decayed gradually with increasing lag, indicating higher-order dependencies beyond adjacent ISIs (Figure 6d, blue).

Critically, we tested whether these ISI correlations reflected intrinsic structure of the saccade generator or were instead due to the slow rate fluctuations. To this end, we applied the time-rescaling transformation, expressing each ISI in units of the integrated intensity of the windowed rate function. After normalization, the serial correlation fell to the level expected under the null band from lag 2 onward, with only a small positive residual remaining at lag 1 (Figure 6d, green). Thus, the higher-order dependencies in the raw ISI sequence were largely due to the slow fluctuations in saccade rate, rather than reflecting intrinsic temporal structure of the saccade generator itself. In other words, once rate fluctuations were accounted for, successive ISIs were approximately consistent with a renewal process, i.e., each ISI primarily depended on the current rate, not on preceding intervals.

Together, these results show that saccade rates within long trials are non-stationary and exhibit slow, scale-free fluctuations that are not merely due to noisy ISI variation. This non-stationarity, specifically the slow rate fluctuation, generates temporal structure in the ISI sequence.

## Discussion

We investigated saccade timing during free viewing in human, macaque, and marmoset. Across the three primate species, saccade timing was well described by a locally rhythmic gamma renewal process with a fluctuating rate. Saccade rates varied widely across trials and fluctuated slowly within long trials, suggesting that the averaging over temporally local regimes creates the empirical heavy-tailed ISI distributions. Indeed, the empirical ISI distributions became more symmetric and narrower after normalizing for slow rate fluctuations. When we introduced empirically observed rate fluctuations into a simulated gamma renewal process with a high shape parameter and thereby high temporally local rhythmicity, this reproduced the empirical heavy-tailed ISIs; notably, this also strongly attenuated previously used rhythmicity metrics based on spectral peaks. Therefore, we introduced the RCRI that is nearly immune to slow rate fluctuations. RCRI values varied across tasks and species, and this variability was largely explained by a genuine dependence of RCRI on saccade rate. Saccade-rate binning revealed that an increasing rate monotonically increases RCRI, for all three primate species. Finally, ISIs showed long-range serial correlations, which collapsed when ISIs were normalized for slow rate fluctuations. Thus, the dominant higher-order structure arose from slow rate fluctuations rather than an intrinsic multi-interval dependence of the rhythmic generator. In summary, heavy-tailed ISI distributions and weak spectral peaks are due to slow rate fluctuations, and normalizing for those fluctuations reveals a robust temporally local rhythmic generator.

Several limitations to our findings should be noted. First, for the macaques and the marmoset, the sample sizes were two and one, respectively, such that inference was limited to the samples rather than applying to the populations^13^. Any comparison between macaques and the marmoset, or between either species and humans, would likewise have been limited to the respective samples, thereby precluding meaningful statistical comparisons between the respective species’ populations. Second, our observations might be specific to the employed stimulus set and viewing regimes, i.e., the free viewing of static dot arrays and natural images, and of relatively slow and smooth movies. For other stimuli with more pronounced temporal structure, or for other behavioral tasks, saccade statistics, including higher-order dependencies, may differ. Third, although the deep learning-based saccade detection is designed to be highly sensitive, rhythmicity and serial-correlation estimates can, in principle, be influenced by systematic detection biases, particularly for small-amplitude saccades and during free viewing where other eye-movement components such as smooth pursuits are present. Yet, our sensitivity analysis suggests that such effects are small to negligible. Finally, our modelling framework is intentionally descriptive and based on behavioral data. It constrains the structure of the generative process but does not identify the underlying neuronal mechanism, motivating future work that links rate modulation and local rhythmicity to circuit-level dynamics.

The temporal structure of saccade sequences has a long history of empirical characterization and theoretical debate. Early work established that successive saccades cannot generally be initiated arbitrarily close in time. This post-saccadic refractory period imposes an apparent hard floor on the ISI, on the order of 100–200 ms ^14,15^ dictating the primary mode of the ISI distribution. Mechanistically, it was suggested by Carpenter and colleagues that this mode reflects the time for a threshold to be reached by a linearly accumulating decision signal with a stochastically varying rate, producing characteristically right-skewed reaction times (or ISI distributions) as a natural consequence of this variability^4,16^. Sumner and colleagues extended this view within neuronal field models to show that competitive fixation-saccade dynamics further shape the tail^17^. Together, these frameworks converge on the view that the heavy-tailed ISI distribution is primarily a product of stochastic accumulation-to-threshold dynamics gated by a post-saccadic recovery period. Even when modelling how saccade generation is interrupted by sensory transients^18^, successive saccade generation was an outcome of repetitive rise-to-threshold waves that could be reset by exogenous stimuli^19^, again utilizing the concept of saccade-system rhythmicity.

The most direct empirical evidence that saccade timing can nonetheless express a genuine local rhythmic component comes from the work of Otero-Millan and colleagues. Examining ISI distributions across saccade amplitude classes during both fixation and free exploration, Otero-Millan et al. ^20^ showed that the statistics of successive events are consistent with a single common saccadic generator whose inter-event distributions share the same ex-Gaussian form regardless of saccade amplitude, with a mean of the Gaussian component near 150 ms and a heavy tail reflecting occasional prolonged intervals. The case for an intrinsically rhythmic generator becomes compelling in patients with square-wave jerks, spontaneous conjugate saccadic intrusions that interrupt fixation in paired saccades away from and back to a fixation location. Otero-Millan et al. ^21,22^ documented that the interval between these paired saccades converges tightly on approximately 200 ms, consistent with square waves observed in healthy individuals^19,23^. Rather than being merely an effect of pathology, these near-constant inter-saccadic periods could be understood as revealing, in unusually clean form, the local rhythmic kernel that the oculomotor system’s intrinsic recovery dynamics impose. This kernel might normally be obscured in healthy free viewing by rate fluctuations, higher cognitive influences, or other sources of variability. This directly supports our finding that a robust rhythmic component is present in healthy primate saccade sequences once slow rate fluctuations are accounted for.

Despite this rich tradition of characterizing saccade timing, none of these accounts set out to quantify saccade rhythmicity as such. To our knowledge, the only study that has considered this question directly is Amit et al., (2017), who explicitly asked whether free-viewing saccade sequences are better described by a self-paced generator or by a fixed-frequency oscillator, i.e., a central pattern generator. To answer this question, Amit et al. quantified spectral modulation strength, complemented by point-process analyses of autocorrelograms and hazard functions, and used ISI shuffling to test for dependencies beyond the immediately preceding saccade. In their data, hazard functions were dominated by post-saccadic inhibition without sustained periodic modulation, autocorrelograms showed at most short-lived damped structure, and shuffling left the oscillatory-modulation index largely unchanged. They concluded that a 3–6 Hz spectral peak is parsimoniously explained by first-order dynamics such as a post-saccadic refractory period, without invoking an autonomous pacemaker.

Our results extend this framework by showing that its core diagnostics can be confounded when saccade rate is non-stationary. Amit et al.’s generative models assume a constant saccade rate, and the ISI shuffle test is naturally interpreted in this stationary setting. In our data, saccade rates show strong slow fluctuations, so the empirical ISI distribution pools intervals sampled at different rates. Rate fluctuations can therefore affect global timing statistics, masking spectral signatures typically attributed to rhythmicity. The same logic applies to the hazard function: locally present periodic modulation can be washed out by the mixture, yielding the flat or monotonically rising profiles that Amit et al. interpreted as evidence against an oscillatory process. Our simulations confirm that shuffling leaves peak power essentially unchanged not only for stationary renewal data but also for non-stationary, rate-modulated data (Figure 3e), so the absence of an effect of shuffling does not uniquely support the inference that only first-order dependencies are present. By contrast, the RCRI is shuffle-invariant for stationary renewal data, but is disrupted by shuffling for non-stationary sequences (Figure 3f), because shuffling breaks the alignment between successive intervals and the slowly varying rate regime that generated them. These results do not contradict Amit et al.’s conclusion that saccade timing is poorly captured by a rigid, continuously phase-advancing oscillator, but extend their analysis to the regime that dominates natural viewing. When rate fluctuations are accounted for, a temporally local rhythmic component becomes evident, while most of the long-range ISI structure is explained by the slow rate fluctuation.

Our descriptive model decomposes saccade timing into a locally rhythmic gamma renewal process and slow rate fluctuations, thereby raising the question of what neuronal architecture could implement this decomposition. A parsimonious candidate is a drift-to-threshold mechanism. In this framework, after each post-saccadic reset, a decision variable accumulates toward a fixed threshold. At threshold crossing, a saccade is triggered, followed by a refractory period before the next build-up begins. Each cycle is analogous to one period of an oscillator, but the frequency is not fixed, it varies with the drift rate and the threshold level. This mechanism directly explains our finding that RCRI increases with mean saccade rate: If the noise of the accumulation process is approximately stable, then increasing the mean drift compresses the time-to-threshold distribution and reduces relative dispersion, producing a more rhythmic local sequence for higher saccade rates (Figure 7).

**Figure 7.**
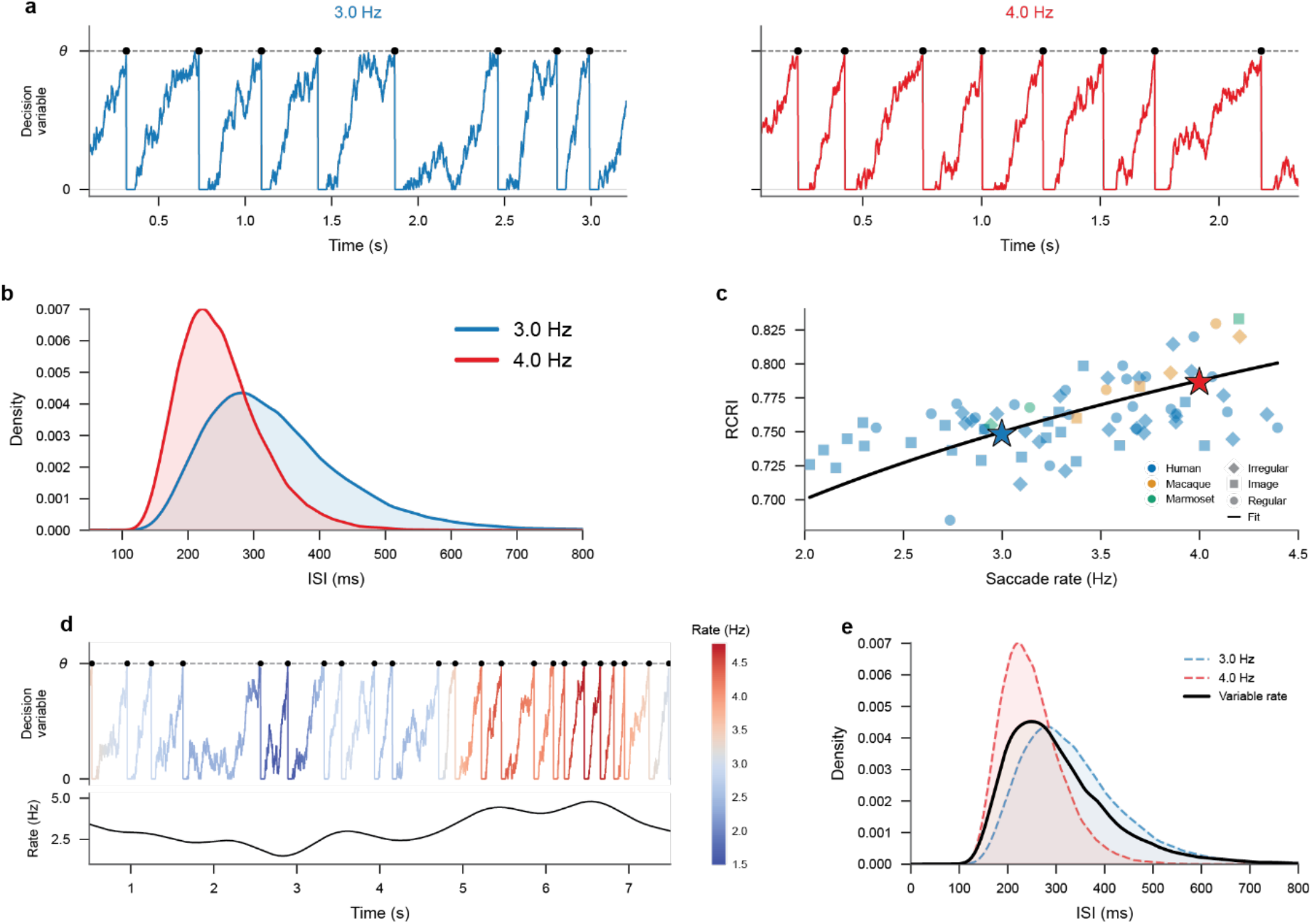
A drift-to-threshold model predicts the observed coupling between saccade rate and rhythmicity. **(a)** Example accumulator traces from a drift-to-threshold model, where a decision variable accumulates with constant drift plus Gaussian noise toward a fixed threshold (dashed line), and upon threshold crossing triggers a saccade. When the model is run with different drift rates, it can produce saccade rates of either 3.0 Hz (blue) or 4.0 Hz (red). **(b)** ISI distributions for the two example rates. Higher drift yields a narrower ISI distribution, corresponding to higher rhythmicity. **(c)** Replotting of the data from Figure 4c, superimposing as black line the rate–RCRI relationship predicted by the drift-to-threshold model. There was only a single free parameter, the Gaussian noise, which was fitted to the empirical data. Stars indicate the two example rates from panels a and b. The model captures the monotonic increase of RCRI with saccade rate observed across species and conditions. **(d)** Accumulator trace driven by a slowly fluctuating rate signal, with color indicating instantaneous rate. The rate time course (bottom) varies smoothly between 1.5 and 4.5 Hz, producing epochs of faster and slower saccade generation. **(e)** ISI distribution from the variable-rate simulation (black) compared to the two constant-rate ISI distributions from panel b (dashed). The variable-rate distribution emerges as a mixture of the underlying constant-rate distributions, demonstrating how slow rate fluctuations broaden the overall ISI distribution.

This mechanistic explanation has strong neurophysiological grounding. In the frontal eye fields (FEF), movement-related neurons reach a relatively fixed discharge level at saccade onset, with latency variability explained by variability in the growth rate toward threshold^24^. In addition, reversible inactivation of the FEF, through cooling, alters the timing of subsequent saccades after sensory-driven interruption^25^. Analogous threshold-crossing dynamics have been reported in the superior colliculus^26^. Downstream, the brainstem burst generator provides an explicit gating and refractory mechanism: omnipause neurons tonically inhibit premotor burst neurons during fixation and pause around saccades, naturally implementing a post-saccadic reset epoch and separating fast, local trigger timing from slower control signals that modulate readiness. Together, these circuits map onto our behavioral decomposition of a locally rhythmic threshold-and-reset generator whose instantaneous rate is set by external drive.

The classic Linear-Approach-to-Threshold-with-Ergodic-Rate (LATER) framework assumes that the drift rate is drawn independently on each saccade cycle (or trial, in the classic LATER model), predicting no long-range serial structure in ISIs beyond the refractory period. The slow non-stationarity we observe violates this assumption: the rate does not reset independently after each saccade but fluctuates slowly, such that successive cycles share a similar local rate context. This can be thought of as the signature of a system operating under slowly drifting control variables such as baseline excitability, tonic fixation drive, and neuromodulatory state, all of which change on timescales longer than a single ISI. Changes in these control variables might well correspond to different cognitive states, like levels of arousal, exploratory drive, reward expectancy and resulting motivation^27–30^.

Finally, this framework makes concrete predictions about how rate and local rhythmicity can dissociate. In drift-to-threshold terms, mean rate varies via the mean drift or the threshold level, whereas rhythmicity at matched rate is determined by the variability in drift, reset depth, or refractory gating. These model parameters might in future studies be linked to neuronal mechanisms through perturbations at various nodes of the saccade control and generation system. At the neuronal level, saccade generation and attentional control are tightly intertwined. Thereby, a deeper understanding of the neuronal sources of saccade rhythmicity will also provide a link to the rich literature on rhythmicity in attentional and perceptual processes ^7,31–33^.

## Acknowledgements

The authors would like to thank Ece Nur Uygur for help with the human data acquisition. This work was supported by DFG (FOR 1847 FR2557/2-1, FR2557/7-1, TH 425/17-1).

## Data, Materials, and Software Availability

All analysis code needed to reproduce the figures and statistics is available at https://github.com/timnaher/saccade_rhythmicity. Eye-movement data are available from the corresponding author upon reasonable request (Tim Näher). The natural image shown in Figure 1c is from the McGill Calibrated Colour Image Database (Olmos and Kingdom, 2004; http://tabby.vision.mcgill.ca) and is not redistributed with the code. The movie used in the free-viewing MOVIE task is the publicly available YouTube video 4K Virtual Hike Near River through the Forest - Baker River Trail and Chain Lake Trail, cited by title only, and is not redistributed with this article for licensing reasons.

## Author contributions

T.N.: Conceptualization, Data curation, Formal analysis, Investigation, Methodology, Resources, Software, Visualization, Writing – original draft, Writing, review & editing. Y.Z.: Conceptualization, Formal analysis, Data curation, Investigation, Methodology, Resources, Software, Visualization, Writing – original draft, Writing, review & editing. S.S.: Investigation, Writing, review & editing. M.P.: Investigation, Writing, review & editing. L.B.: Conceptualization, Writing, review & editing. P.T.: Supervision, Funding acquisition, Writing, review & editing. Z.H.: Conceptualization, Methodology, Writing – original draft, Writing, review & editing. P.F.: Conceptualization, Supervision, Funding acquisition, Writing – original draft, Writing, review & editing.

## Declaration of generative AI and AI-assisted technologies in the manuscript preparation process

During the preparation of this work, the authors used Claude Opus and ChatGPT to aid with writing the manuscript. After using these tools, the authors reviewed and edited the content as needed and take full responsibility for the content of the published article.

