## Supplementary Methods for "Primate Saccade Rhythmicity"

### Participants and animals

Twenty-five adult human participants (12 males, 13 females), recruited from the public, participated in the current study. We excluded three participants (all female) due to insufficient numbers of trials. All human participants had normal vision and provided informed consent before participation.

Two adult rhesus macaques (*Macaca mulatta*, denoted C and G in the main text, both 19 years old) participated in this study. Both animals were implanted with a headpost. Monkey G had a recording chamber implanted for addressing other scientific questions

One adult marmoset (*Callithrix jacchus,* 6 years old) participated in the study. It was implanted with a headpost.

### Ethics statement

***Humans***

The study was approved by the local ethics committee (Ethikkommission des Fachbereichs Medizin der Goethe-Universität). Participants gave written consent before participating in the study.

***Macaques***

All experimental protocols were approved by the responsible authority (Regierungspräsidium Darmstadt) and complied with the applicable German and European legislation, including EU Directive 2010/63/EU on the protection of animals used for scientific purposes.

***Marmoset***

All experimental protocols were approved by the responsible authority (Regierungspräsidium Tübingen) and complied with the applicable German and European legislation, including EU Directive 2010/63/EU on the protection of animals used for scientific purposes.

### Tasks

Except for the MOVIE task, performed only by human participants, all species performed the same REGULAR, IRREGULAR and IMAGE tasks with only minor differences in the reward procedure (auditory feedback for humans versus juice delivery for macaques and marmoset). Figure 1 illustrates the stimuli used in the REGULAR, IRREGULAR and IMAGE tasks.

***Macaque tasks***

All recording were performed during head fixation. Macaques were recorded in two types of sessions. Each type of session contained a specific set of tasks.

Session type 1 consisted of the REGULAR and IRREGULAR tasks. Each trial in both tasks began with the animal fixating at the center of the screen on a fixation dot (diameter 0.1 dva, i.e. degrees of visual angle). After fixation for 800 to 1000 ms, the fixation dot was turned off, followed by a ≈100 ms gap. The search array was then presented. The array consisted of white filled dots (diameter 0.3 dva) that reflected potential saccade targets (see Figure 1 for an illustration). Within the search array, 20% of the presented dots were real targets. The animal's task was to sequentially look at the dots to find any one of the real targets (foraging). During the foraging, if the animal's gaze landed inside a fixation window around a real target (radius 1.0 dva, not visible to the monkey), that target was replaced by an image of grapes, and a small juice reward was provided. If the monkey kept fixation on the grapes, multiple pulses of reward were provided every 200 ms until a maximum window of 2000 ms had passed. This was followed by an inter-trial interval of 500 ms and the start of a new trial with the presentation of a new fixation point. If the maximum search time passed (5000 ms), and none of the real targets had been foveated, the trial was terminated without a reward, followed by the same inter-trial interval and the start of a new trial. In the REGULAR task, the dots in the search array were spaced as if they were the centers of an invisible hexagonal tiling, with 4.5 dva between adjacent dots. In the IRREGULAR task, the search array contained the same number of dots as in a REGULAR trial, but the positions of the dots were randomly redistributed across the full screen to break any spatial regularity.

Session type 2 consisted of only one task, the IMAGE task, and began with the animal fixating at the central fixation dot. This pre-search fixation for the IMAGE task was identical to the REGULAR or IRREGULAR tasks. After fixation for 800 to 1000 ms, the fixation dot was turned off, and after a ~100 ms gap, it was replaced by a natural image in color (16 * 21 dva, McGill Calibrated Colour Image Database, http://tabby.vision.mcgill.ca). From the moment the fixation dot was turned off, the animal was free to explore the image. After a random delay of 100 ms to 1900 ms, a small white dot (either 0.1 dva radius or 1.7 dva radius, randomly interleaved), the target, appeared at a random location within the image. We used two different radii for a comparison that is not relevant to the current study. The monkey's task was to locate this target within a period of 5000 ms. If the target dot was found, the reward procedure as described for session type 1 followed.

***Marmoset tasks***

All recording were performed during head fixation. The marmoset tasks closely mirrored the macaque version, with parameter adjustments to accommodate species-specific behavioral and attentional constraints. The key differences are described below.

All three tasks (REGULAR, IRREGULAR, and IMAGE) were presented within a single session type, in random order. Each trial began with fixation on a central red dot (diameter 0.4 dva) which the animal had to acquire within 200 ms and hold for up to 600 ms. The fixation dot was then turned off and followed by a ~100 ms gap. In the REGULAR and IRREGULAR task, the search array was then presented. Array composition, target proportion (20%), dot spacing rules, and maximum search time (5000 ms) were identical to the macaque version.

The fixation window around real targets was enlarged (2 x 2 dva, not visible to the monkey) to reduce frustration and encourage longer working times. Upon successful target fixation, the target was replaced by an image of a marmoset face (3 × 4 dva), and a small amount of marshmallow juice was delivered. If the monkey kept fixation on the marmoset face, short pulses of reward were provided manually until trial end.

For the IMAGE task, stimuli (16 x 21 dva) were drawn from the same database. Only a single target size was used (diameter 0.3 dva, white). The monkey's task was to locate this target within a period of 5000 ms. If the target dot was found, multiple pulses of reward were manually provided until a maximum window of 3000 ms had passed.

Note that due to a bug in the stimulation script, the end-of-trial marker was missing in the marmoset data. We therefore decided on a fixed trial length of 4 s, in which the marmoset found most of the rewards.

***Human tasks***

All recordings were performed with participants' heads stabilized using a chinrest. Human participants were recorded in two types of sessions. Each type of session contained a specific set of tasks. Session type 1 consisted of the REGULAR, IRREGULAR, and IMAGE tasks. These tasks for human participants were identical to the respective tasks for macaques, except for the following differences: (1) The pre-search fixation duration was 2500 to 3500 ms; (2) the post-search fixation duration was also 2500 to 3500 ms and did not contain a repetitive reward; (3) in all tasks, successful trial completion was indicated by a short audio feedback. If the maximum search time passed and no target was found, the trial was terminated. Inter-trial-intervals were 2000 ms.

Session type 2 included the REGULAR and IMAGE tasks, as well as the free-viewing MOVIE task. It was identical to session type 1, except for the following differences: (1) The REGULAR task in session type 2 contained three different spacings of the grid (hexagon radius of 2.3, 4.5, and 8.5 dva); (2) the IMAGE task remained unchanged from session type 1 except for a different set of images being used.

The free-viewing MOVIE task (performed exclusively by human participants) consisted of a 10-minute naturalistic first-person-view video. The video contained minimal cuts, such that uninterrupted epochs lasted several minutes. Furthermore, the camera trajectory was smooth, minimizing transients due to camera movement. The movie is in color, was downloaded from YouTube (title 4K Virtual Hike Near River through the Forest - Baker River Trail & Chain Lake Trail) and can be obtained from the authors upon request. We chose this movie, because it provides naturalistic visual content, motivates subjects to sustained saccadic exploration of the continually renewing input, and largely avoids transients. Participants were instructed to keep their head still on the chinrest and simply watch the movie without performing a specific task.

The human data were collected across one or two sessions, with some participants attending only the first session and others attending both. The first session included a brief training block to ensure participants gained experience with the online eye-tracking control and to fully understand the task. A given session encompassed up to six blocks with the REGULAR, IRREGULAR, and IMAGE tasks randomly interleaved. Each block consisted of 60 trials, 20 of each task. Participants worked on one block until 60 correct trials (20 of each condition) were achieved. Participants were encouraged to complete all six blocks; however, the total number completed was contingent upon each participant's level of fatigue. The experimental design sought to prioritize participant comfort and data accuracy over the volume of completed blocks. The second session used the same approach, but for session type 2.

### Eye Movement Recording

All visual stimulation was controlled by custom software (https://github.com/esi-neuroscience/ARCADE) and presented on an LCD monitor. For macaques, the monitor was an LG 32GK850G-B with a refresh rate of 143.9 Hz. For marmoset monkeys, the monitor was a 10 Inch Beetronics with a refresh rate of 60 Hz. For human participants, the monitor was a ViewPixx (VPixx Technologies Inc) with a refresh rate of 120 Hz. In humans and macaques, we recorded binocular gaze data at 500 Hz using an EyeLink-1000 system (SR Research Ltd.). In the marmoset monkey, we recorded monocular gaze data at 120 Hz using an ISCAN ETL-200 infrared eye tracking system. All gaze data were up-sampled to 1 kHz to achieve a common sampling rate for all further analyses. Real-time gaze data from one eye was utilized for online behavioral control such as the pre- and post-search fixation periods.

### Saccade detection

We used a convolutional neural network (CNN) for saccade detection[1]. We trained separate networks for humans, macaques, and the marmoset on randomly selected trials from each species. First, we labeled saccades and fixations as separate classes in the data using a custom GUI. After epoching the data into trials, we replaced blinks with NaNs. Subsequently, we used those preprocessed data to train the CNN, then detected saccades for each eye separately. We only accepted saccades if they occurred in both eyes simultaneously. For all detected saccades, we enforced a minimum duration of 6 ms and a minimum separation between two subsequent saccades of 20 ms.

### Estimation of the gamma shape parameter

To quantify the rhythmicity of the inter-saccadic intervals (ISIs) independently of the overall saccade rate, we modeled the ISIs within each trial that contained at least two ISIs using a gamma renewal process. Specifically, for trial $t$, the $n_t$ observed ISIs, $I_{t,1}, \cdots, I_{t,n_t}$, were assumed to be independent draws from a gamma distribution with a common shape parameter $k$ and a trial-specific scale parameter $\theta_t$. We parameterized the scale as

$$\theta_t = \frac{1}{k\lambda_t}$$

, where $\lambda_t$ denotes the prevailing saccade rate in trial $t$. The shape parameter $k$ is common across trials and determines the rhythmicity of the ISIs: for a gamma distribution, the coefficient of variation is $1 / \sqrt{k}$, and the skewness is $2 / \sqrt{k}$. Thus, larger values of $k$ correspond to more rhythmic ISIs, with $k = 1$ corresponding to the exponential ISI distribution of a Poisson process.

We sought to estimate $k$ without introducing trial-specific estimates of the saccade rate. This is important because estimating the rate from the trial duration and the number of saccades can be sensitive to whether a saccade happens to fall just inside or outside the trial boundaries. We therefore based the analysis exclusively on the observed ISIs. Moreover, estimating the rate from the same ISIs that are subsequently normalized creates a dependence between the normalization and the observations whose variability is being assessed.

For each trial, we therefore calculated the total duration represented by the observed ISIs,

$$S_t = \sum_{i=1}^{n_t} I_{t,i}$$

and expressed each ISI as a proportion of this total,

$$P_{t,i} = \frac{I_{t,i}}{S_t}$$

Thus, $P_{t,i}$ indicates what fraction of the total interval time of the trial is occupied by ISI $i$. The proportions are positive and sum to one. Importantly, dividing all ISIs by their trial-specific sum removes the overall timescale of the trial, and hence the prevailing saccade rate, while retaining the relative temporal structure of the ISIs. For example, multiplying all ISIs of a trial by the same factor changes the overall rate but leaves all proportions unchanged.

The gamma distribution provides a particularly useful property for this normalization. If the absolute ISIs follow a gamma distribution with shape $k$, then their proportions of the total follow a Dirichlet distribution with the very same $k$. More specifically, normalizing the ISIs by their trial-wise sum removes their common scale and transforms their joint distribution into a Dirichlet distribution. In mathematical notation,

$$\left(P_{t,1}, \cdots, P_{t,n_t}\right) \sim Dirichlet(k, \dots k)$$

Here, $Dirichlet$ denotes the probability distribution of a set of positive proportions that sum to one, and $(k, \dots k)$ simply means that the same parameter $k$ is assigned to each of the $n_t$ proportions. Because the gamma-to-Dirichlet transformation preserves the shape parameter $k$, $k$ can be estimated from the rate-independent proportions and then interpreted directly as the shape parameter of the underlying gamma ISI distribution. The distribution of the proportions depends on $k$, but not on the trial-specific scale $\theta_t$ or rate $\lambda_t$. Thus, the rate is eliminated analytically rather than estimated and subsequently used to rescale the ISIs.

We estimated the common shape parameter $k$ by maximum likelihood from the Dirichlet distributions of the interval proportions across all trials of each participant. Specifically, the likelihood for each trial that contained at least two ISIs was determined by the observed proportions and their number $n_t$, and the log-likelihoods were summed across trials. The resulting likelihood depends on $k$ as the only free parameter, which we optimized numerically for $k > 0$.

Goodness of fit was assessed on the interval proportions, and the distribution under the model is known. Each proportion follows a beta distribution determined by the shape parameter and the number of intervals in that trial. We therefore expressed each observed proportion as its percentile under that distribution. If the model holds, these percentiles are uniformly distributed regardless of trial length, so trials of different lengths can be pooled. Goodness of fit was then assessed with percentile–percentile plots against the uniform distribution, and formally with a one-sample Kolmogorov–Smirnov (KS) test. A non-significant statistic indicates that the data are consistent with the model.

For each subject (humans, macaques, and marmoset), we extracted all valid ISIs. For model fitting and visualization, all analyses used ISIs within 100–600 ms range, thereby matching the range that we used for histogram-based comparisons.

***Leave-one-out normalized ISIs for visualization***

To visualize the distribution of rate-corrected ISIs while avoiding a contribution of each ISI to its own normalization factor, we normalized each ISI by the mean of all other ISIs in the same trial (for all trials that contained at least two ISIs). Specifically, for trial (t) and ISI (i), we calculated

$$R_{t,i}^{(-i)} = \frac{I_{t,i}}{\frac{1}{n_t - 1}\sum_{j/=i} I_{t,j}} = \frac{(n_t - 1)I_{t,i}}{S_t - I_{t,i}}$$

where $n_t$ is the number of ISIs in the trial, and $S_t = \Sigma_{j=1}^{n_t} I_{t,j}$ is the total duration of all ISIs in that trial. Thus, each ISI is normalized by the mean of the remaining ISIs from the same trial, without contributing to its own normalization. A value of 1 indicates that the ISI equals the mean of the other ISIs, whereas values above or below 1 indicate longer or shorter ISIs, respectively.

This normalization is directly related to the rate-independent Dirichlet formulation described above. Defining the interval proportion $P_{t,i} = I_{t,i} / S$, the leave-one-out normalized ISI can be written as

$$R_{t,i}^{(-i)} = (n_t - 1)\frac{P_{t,i}}{1 - P_{t,i}}$$

Importantly, estimation of $k$ was based on the joint Dirichlet likelihood of the interval proportions; the leave-one-out normalized ISIs were used only for visualization.

### Rate-corrected rhythmicity index

To quantify local saccade rhythmicity while minimizing sensitivity to slow saccade-rate fluctuations, we adopted a metric previously introduced to quantify the neuronal discharge variability based on the inter-spike-interval coefficient of variation ($C_V$), i.e., the ratio of the standard deviation to the mean. Note that in the case of a Poisson process, the inter-spike intervals have a $C_V$ of one, whereas the spike counts have a Fano factor of one, with the Fano factor being the ratio of variance to mean. Furthermore, the $C_V$ of inter-spike intervals is immune to scale changes, i.e., whether intervals are expressed e.g. in millisecond or seconds. Therefore, the $C_V$ has been the standard metric to quantify the variability of inter-spike intervals [2]. Yet, when the $C_V$ is calculated across many ISIs of a temporally extended recording, while the spike rate is changing, it will not provide an accurate estimate[2]. Therefore, a previous study introduced $C_{V2}$, which avoids sensitivity to changing rates by comparing only adjacent intervals[2]. Specifically, $C_{V2}$ is defined as

$$C_{V2}(t,i) = \frac{2\left|I_{t,i+1} - I_{t,i}\right|}{I_{t,i+1} + I_{t_1 i}}$$

The overall local variation was computed as the average $C_{V2}$ over all intervals. While $C_{V2}$ provides a rate-insensitive metric of interval variability, we aimed at a rate-insensitive metric of rhythmicity, and therefore defined the rate-corrected rhythmicity index as

$$RCRI = \frac{1}{1 + C_{V2}}$$

For a Poisson process, the expected value of $C_{V2}$ is 1, yielding $RCRI = 0.5$. Values of $RCRI > 0.5$ therefore indicate greater local rhythmicity than expected from a Poisson process. RCRI inherits the scale invariance of $C_{V2}$: $C_{V2}$ is a ratio of differences to sums of consecutive ISIs, such that multiplying all intervals by a constant leaves the index unchanged.

A sensitivity analysis of how RCRI depends on analysis-parameter choices is shown in *SI appendix*, Fig. S1A.

### Trial-wise saccade rates

For all further analyses of the trial-based conditions (i.e. REGULAR, IRREGULAR, IMAGE, but not MOVIE), only trials with a minimum duration of 1 s were included.

A central aspect of the analyses was concerned with saccade rates. For the trial-based conditions, we computed the mean saccade rate for each trial and used this to quantify between-trial variability in saccade rate. Saccade rate was computed as the total number of detected saccades within the trial divided by the trial duration in seconds, after excluding the initial 100 ms post stimulus onset, because saccade rates are known to show a transient decrease after fixation or stimulus onset[3–6]. Trial-wise saccade rates were used for all analyses relating saccade rate to the rate-corrected rhythmicity index (RCRI, see below), including rate stratification and regression analyses.

### Stationary versus non-stationary processes and spectral measures

To isolate the effects of slow rate fluctuations on rhythmicity metrics and spectral measures, we simulated event sequences using stationary and non-stationary gamma renewal processes that were matched in local regularity but differed in their rate dynamics.

In both simulations, saccade times were generated as a gamma renewal process, in which successive ISIs were drawn independently from a gamma distribution. Local regularity was held constant by fixing the coefficient of variation of the ISI distribution to $C_V = 0.15$, corresponding to a highly rhythmic, oscillator-like, generator.

In the stationary process, the mean ISI was fixed by setting a constant rate equal to the mean of an empirical rate time course derived from the MOVIE task, resulting in a time-invariant ISI distribution.

In the non-stationary process, the mean ISI varied over time according to that same empirical rate time course from the MOVIE task. At each step, the rate of the gamma process was set to the inverse of the current empirical rate, while the shape of the distribution, and thus local regularity, was kept constant. As a result, ISIs were locally regular but globally heterogeneous due to slow rate fluctuations present in the empirical rate dynamics.

***Spectral analysis of event trains***

To quantify rhythmicity with spectral measures, simulated event times were discretized into a binary event train at 1 ms resolution (sampling frequency 1000 Hz). Power spectral density (PSD) estimates were computed using Welch's method (segment length 2048 samples, constant detrending, density scaling).

We generated multiple (n = 500) independent realizations of the stationary and non-stationary processes. For each realization, we extracted the peak power in the 2–6 Hz range[7] and the $RCRI$ computed from the simulated ISIs.

### Rate-stratified analysis of saccade rhythmicity

To assess the relationship between saccade rate and rhythmic structure using a nonparametric approach, we performed a rate-stratified analysis within each species separately. For each trial, mean saccade rate was computed as defined earlier.

Within each species, trials were ranked by mean saccade rate and assigned to tertiles (low, medium, high) using quantile binning. This procedure controlled for differences in saccade rate across species and allowed rate-dependent effects to be examined within comparable relative-rate ranges.

For each species and rate bin, saccade autocorrelation functions were computed within the bin and then averaged over subjects. Additionally, ISI distributions were obtained by pooling ISIs across trials within the bin. For each trial, the RCRI was obtained by averaging $C_{V2}$ across all consecutive ISI pairs within that trial and then applying RCRI = 1/(1 + mean $C_{V2}$); the resulting trial-wise RCRI values were grouped by rate bin.

For humans, pairwise comparisons of rhythmicity between rate bins were performed using Welch's unequal-variance t-tests. For macaques and marmoset, where trials were pooled across individuals[8], pairwise comparisons were performed using two-sided

Wilcoxon rank-sum tests. All statistical tests were conducted within species, and significance was assessed at $\alpha = 0.05$.

### Within-trial slow rate fluctuations and induced higher-order dependencies

***Windowed saccade-rate estimates***

Within each MOVIE trial, saccade onset times were binned into non-overlapping windows of 3 s, yielding a within-trial rate time series.

Blinks were identified as time segments in which eye-position samples contained NaNs. For analyses based on ISIs, any ISI which overlapped a blink segment was excluded For each window, saccade rate was computed as the number of saccade onsets divided by the non-blink duration of that window, so that blink time was excluded from both the numerator and the denominator. Windows in which more than half of the samples were missing were excluded from the rate time series.

For each trial, we computed the lagged autocorrelation of the windowed saccade-rate time series for lags up to 20. Autocorrelation functions were then averaged over trials.

To characterize the frequency content of within-trial rate fluctuations, we estimated the power spectral density of the windowed rate series using Welch's method. Because rate was sampled once per 3 s window, the effective sampling frequency was 1/3 Hz.

***Monte Carlo null model for rate autocorrelation***

To test rate autocorrelations against a memoryless baseline, we constructed a per-trial surrogate under a homogeneous Poisson null while preserving each trial's total saccade count. For each eligible trial with $N_i$ saccades and $B_i$ windows, surrogate window counts were generated via a multinomial draw with uniform bin probability $1/B_i$. Surrogate rates were computed as counts divided by 3 s. The autocorrelations were computed identically to the empirical data.

This procedure was repeated 500 times. For each lag, we derived a 95% null envelope based on the 2.5th–97.5th percentiles.

***ISI serial correlations***

For each trial, we again computed the ISI sequence as the difference between successive saccade onset times (see above). Higher-order temporal dependencies were quantified using lagged serial correlations across the ISI sequence: for $k = 1, \dots, 20$, the correlation between $ISI_n$ and $ISI_{n+k}$ was computed. Correlation functions were averaged across trials.

***Rate-conditioning via integrated intensity transformation***

To assess ISI serial correlations after conditioning on within-trial rate fluctuations, we applied a time-rescaling transformation using the trial's windowed rate estimate. This rescaling allows us to analyze ISI time courses, while removing correlations introduced by rate fluctuations.

1. The windowed rate series was linearly interpolated to obtain a continuous rate function $\lambda(t)$, with $\lambda(t) \geq 0$.
2. For each ISI spanning $[t_n, t_{n+1}]$, we computed the integrated intensity

$$\Lambda_n = \int_{t_n}^{t_{n+1}} \lambda(t)\, dt,$$

using numerical integration with a step size of 0.01 s. Importantly, $\Lambda_n$ gives the duration of the interval $[t_n, t_{n+1}]$ measured in rescaled (intensity) time. This transformation expresses each ISI in units of expected event count, thereby removing correlations induced solely by slow rate fluctuations.

3. Serial correlations were then recomputed on the transformed interval sequence $\{\Lambda_n\}$ using the lag range (1–20).

To establish the baseline expected from the rescaling itself, we generated 100 surrogate saccade trains per trial that carried the same estimated rate fluctuations but no further dependency, and passed each through the identical pipeline. Because the rate is estimated from the same saccades it rescales, the resulting 95% band (2.5th–97.5th percentiles across surrogates), rather than zero, is the reference for the rescaled serial correlations.

### Drift-to-threshold model

To formalize the relationship between saccade rate and rhythmicity, we implemented a drift-to-threshold model (Figure 7; similar to LATER models)[9]. In this model, a decision variable accumulates linearly with drift rate $\mu$ and additive Gaussian noise (diffusion coefficient $\sigma$) toward a fixed threshold $\vartheta$. Upon threshold crossing, a saccade is triggered, and the accumulator resets to zero, followed by a fixed refractory period $\tau$. The resulting inter-saccade intervals follow an inverse-Gaussian distribution with mean $\vartheta/\mu$ and shape $\vartheta^2/\sigma^2$, shifted by $\tau$. We fixed the threshold at $\vartheta = 1$ (arbitrary units) and the refractory period at $\tau = 50$ ms, and fitted the single remaining free parameter σ to the scatter plot of RCRIs as function of empirical rates, as presented in Figure 4, by minimizing the sum of squared errors. The fit was restricted to the 74 participant × condition cells with a saccade rate of at least 2 Hz, spanning 2.02–4.39 Hz, and yielded σ = 0.71 (RMSE = 0.025). For each empirical saccade rate $r$, the corresponding drift was determined analytically as $\mu = \vartheta / (1/r - \tau)$. Predicted RCRI values were computed deterministically via numerical integration over the inverse-Gaussian probability density function on a 500-point grid. To illustrate how natural rate fluctuations affect rhythmicity, we additionally simulated an accumulator driven by a time-varying drift signal with $1/f$ spectrum, producing a continuous mixture of saccade rates between 1.5 and 4.5 Hz.

### Statistical analyses

To assess whether trial-wise saccadic rhythmicity could be explained by saccade rate, we performed species-specific linear regression analyses using ordinary least squares. The dependent variable was trial-wise RCRI, as defined above. Mean saccade rate (Hz) was computed for each trial as defined above.

For humans and macaques, rhythmicity was modelled as a function of mean saccade rate and experimental condition. Condition was included as a categorical predictor with three levels (REGULAR, IRREGULAR, IMAGE).

For humans, additional comparisons of rhythmicity across rate-defined bins were performed using Welch's unequal-variance t-tests. For macaques, where data were pooled across individuals, and for the marmoset, comparisons were performed using two-sided Wilcoxon rank-sum tests.
Unless otherwise stated, all statistical significances were assessed at $\alpha = 0.05$. All analyses were conducted in Python 3.8.8 using NumPy (v1.22.4), pandas (v2.0), SciPy (v1.10.1), and *statsmodels* (v0.14).
