## Supplementary Figure for "Primate Saccade Rhythmicity"

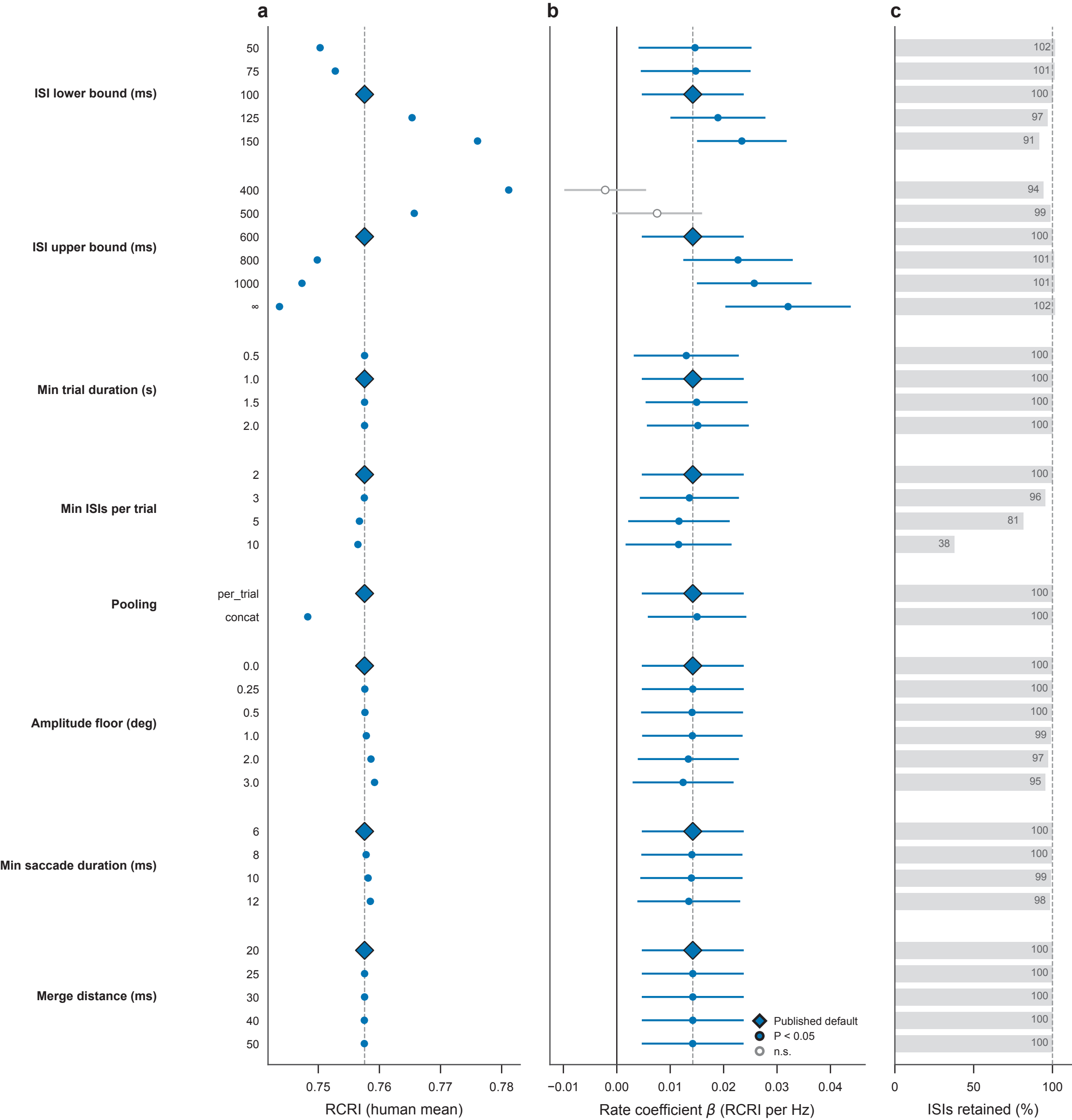

a
b
c
ISI lower bound (ms)
50
75
100
125
150
ISI upper bound (ms)
400
500
600
800
1000
∞
Min trial duration (s)
0.5
1.0
1.5
2.0
Min ISIs per trial
2
3
5
10
Pooling
per_trial
concat
Amplitude floor (deg)
0.0
0.25
0.5
1.0
2.0
3.0
Min saccade duration (ms)
6
8
10
12
Merge distance (ms)
20
25
30
40
50
0.75
0.76
0.77
0.78
RCRI (human mean)
−0.01
0.00
0.01
0.02
0.03
0.04
Rate coefficient β (RCRI per Hz)
Published default
P < 0.05
n.s.
102
101
100
97
91
94
99
100
101
101
102
100
100
100
100
100
96
81
38
100
100
100
100
100
99
97
95
100
100
99
98
100
100
100
100
100
0
50
100
ISIs retained (%)
